# Ultrafast X-Rays Capture Retinal Traversing Conical Intersection in Rhodopsin

**DOI:** 10.64898/2026.07.10.737858

**Authors:** Thomas D. Grant, Suchithranga M.D.C. Perera, Leslie A. Salas-Estrada, C. Swathi K. Menon, Andrey V. Struts, Xiaolin Xu, Steven D. E. Fried, Nipuna Weerasinghe, Udeep Chawla, Roberto Alvarez, Hao Hu, Konstantinos Karpos, Stella Lisova, Reza Nazari, Sahba Zaare, Jesse Coe, Raimund Fromme, Domingo Meza, Abhishek Singharoy, Sarah R. Chamberlain, Stephen Moore, Nadia A. Zatsepin, Fivos Perakis, Sergio Carbajo, Mark S. Hunter, Mengning Liang, Matthew D. Seaberg, Sébastien Boutet, Derek Mendez, Alan Grossfield, Petra Fromme, Richard A. Kirian, Michael F. Brown

## Abstract

G-protein-coupled receptors of the *Rhodopsin* family are crucial medicinal targets, transmitting signals across biomembranes. While light absorption by visual rhodopsin is well studied, its activation via retinal cofactor dynamics remains unclear. Here we use a free-electron laser to show that time-resolved X-ray solution scattering captures the retinal *cis–trans* isomerization as it passes the conical intersection of excited and ground-state photoproduct energy surfaces. Femtosecond-scale nuclear changes occur due to resonant photon absorption, with all-atom simulations revealing ultrafast amino acid movements that initiate transmembrane helix shifts. Ligand-free opsin measurements confirm that light activation is unaffected by non-resonant processes, showing the photonic energy is directly transmitted within the protein. Our method unveils how cofactor dynamics activate rhodopsin, free of constraints from crystal lattice packing or cryotrapping photointermediates.

## Main Text

Photoisomerization of the retinal cofactor of animal and microbial rhodopsins is among the fastest reactions in biology with a high efficiency and ultrafast dynamics ^1,2^. For visual rhodopsin—the G-protein-coupled receptor (GPCR) involved in dim-light perception—optical spectroscopy ^3,4^ has shown the primary rhodopsin photoproduct forms within ∼200 fs ^4–8^. Retinal isomerization underlies visual signaling ^9^, ion conduction, and energy transduction by microbial (archaeal and bacterial) rhodopsins ^10^, and optogenetics applications ^11^. Current structural information is derived largely from crystallographic approaches ^12–17^, NMR spectroscopy ^9^, and cryogenic electron microscopy ^18^ combined with machine-learning ^19^. Light absorption by retinal is followed by excited state decay which takes on average ∼90 fs as indicated by quantum chemical calculations ^1,2^ (1 fs = 10^−15^ s). From ultrafast absorption ^7^ and transient-grating ^8^ measurements of rhodopsin, it is postulated the transition state involves a conical intersection (CI) ^20^ of the excited reactant and ground-state photoproduct energy surfaces, yielding isomerization of retinal from 11-*cis* to all-*trans* in a vibrationally coherent manner (Fig. 1). The theoretical CI point is crucial to explaining the dual requirements of low basal (dark-state) activity together with ultrafast reaction dynamics and high efficiency that enable rhodopsin to detect low light levels in scotopic vision.

**Fig. 1.**
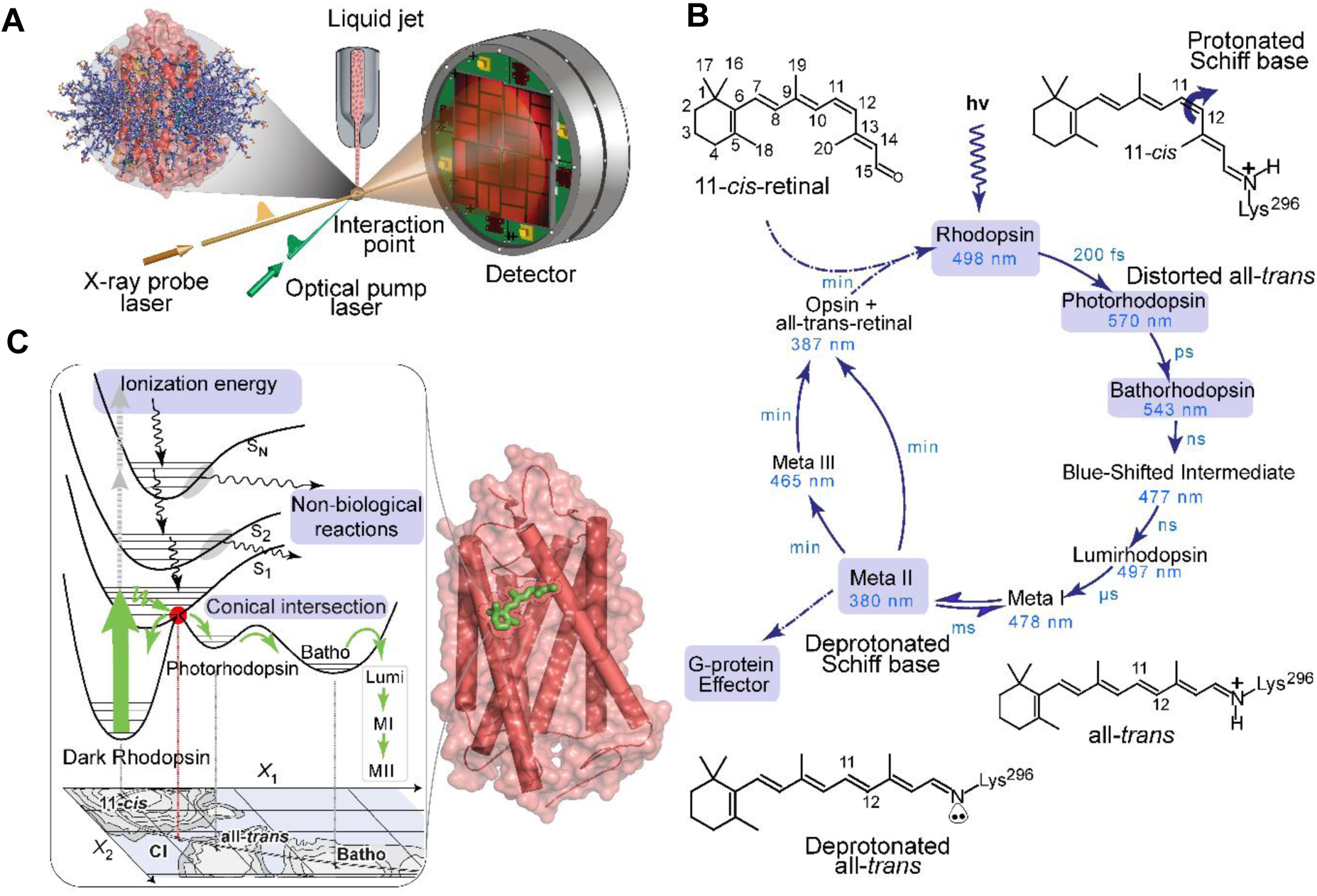
Exploring ultrafast photochemistry of visual rhodopsin with femtosecond X-ray scattering. **(A)** Sample delivery and laser excitation scheme for time-resolved pump-probe studies of rhodopsin solubilized in CHAPS detergent micelles at Linac Coherent Light Source (LCLS). (**B**) Sequence of rhodopsin photointermediates from optical spectroscopy ranging from fs to ms due to photolysis of the bound retinal chromophore. (**C**) Electronic energy states of visual rhodopsin (PDB accession code: 1U19) ^13^ described by potential surfaces of retinal cofactor. Absorption of a green incident photon yields vertical excitation to an excited state, which relaxes to the conical intersection (CI) with the ground-state S0 potential surface ^23^.The protein environment channels the excited-state energy through the CI yielding the biologically relevant pathway. Note the extracellular side (e-side) of rhodopsin is at top and the cytoplasmic side (c-side) at bottom.

With decades of research, the direct structural detection of the sub-picosecond isomerization of retinal in real time—the Holy Grail of visual photochemistry—has eluded researchers until now. Time-resolved crystal structures have recently become available^15^, yet exploring the sub-picosecond dynamics at high temporal resolution remains challenging. Furthermore, confinement in a crystal lattice can alter biochemical pathways even in serial crystallography ^21^. Here we show that time-resolved X-ray solution scattering (TR-XSS) with an X-ray free electron laser (XFEL) (Fig. 1A) detects the ultrafast *cis–trans* isomerization of retinal in real time while passing through the CI point for visual rhodopsin. Fleeting atomic motions during barrier crossings are what defines chemistry, allowing us to observe a protein in action. Our detection method rests on comparing the rhodopsin holoprotein to ligand-free opsin to establish the differences in X-ray scattering that come from resonant light absorption by the retinal chromophore (Fig. 1B). Besides isomerization of retinal (Fig. 1C), ultrafast motions of highly conserved amino acid residues occur near the cofactor binding pocket as shown by our molecular dynamics simulations. By circumventing crystal lattice packing forces, cryotrapping of photointermediates, and radiation damage to crystalline order, our method brings significant advantages to exploring protein dynamics in structural biology.

### Ultrafast dynamics of visual receptors seen by femtosecond X-ray scattering

For rhodopsin, we used ultrafast X-ray scattering to investigate the dynamical processes due to light absorption in the initial stage of vision, before large-scale protein conformational changes occur ^10^. This study marks the first femtosecond time-resolved scattering experiment with a GPCR such as rhodopsin. The ultrafast XFEL at LCLS allowed us to explore how the functional conformational changes of rhodopsin are initiated by dynamic allostery of the retinal ligand. To achieve this goal, we developed methods for time-resolved X-ray scattering using an XFEL source. Our experiments ensured that data collection outpaced radiation damage (diffraction before destruction) by using ultrafast 40-fs XFEL pulses. The protocol encompassed optimizing the sample delivery system, control apoprotein samples, and all-atom molecular dynamics (MD) simulations with an explicit solvent.

Our aim for rhodopsin was to probe the ultrafast nuclear changes (0–10 ps) due to photonic energy absorption that precede activation of the receptor. We hypothesized that photoisomerization of retinal could be detected via X-ray scattering of the protein in a noncrystalline environment ^9^. Accordingly, we acquired TR-XSS data of rhodopsin in non-crystalline environment (CHAPS detergent micelles to follow the protein structural changes near ambient temperature (Fig. 1A). In our pump-probe experiment, an ultrafast pulse of visible light excites rhodopsin molecules flowing in a liquid jet, initially in the dark, nonexcited state. Shortly after the pump excitation with an optical parametric amplifier (OPA) (∼70-fs pulse width), the probe XFEL pulse (40-fs width) scatters X-rays from the rhodopsin molecules onto the detector. Convolving the Gaussian pulse durations yields ∼80 fs as the nominal time resolution. The X-ray scattering profiles were collected versus the delay time after excitation, which ranged from –2 ps to +10 ps ^22^. Use of the LCLS time tool ^22^ allowed us to reduce the ∼300 fs timing uncertainty to ∼50 fs. The averaged laser-off scattering profiles were subtracted from the corresponding laser-on profiles to isolate the signal from the photon absorption (materials and methods) (Fig. 2A, B). For rhodopsin the TR-XSS data showed a significant difference-scattering signal at *q*-values less than ∼0.12 Å^−1^, whereas the higher *q*-range had little discernible signal above the noise (Fig. 2A, 3D). The light-induced negative difference scattering signal of rhodopsin is readily seen in the 1D difference scattering profiles in the low-*q* range (Fig. 2C).

**Fig. 2.**
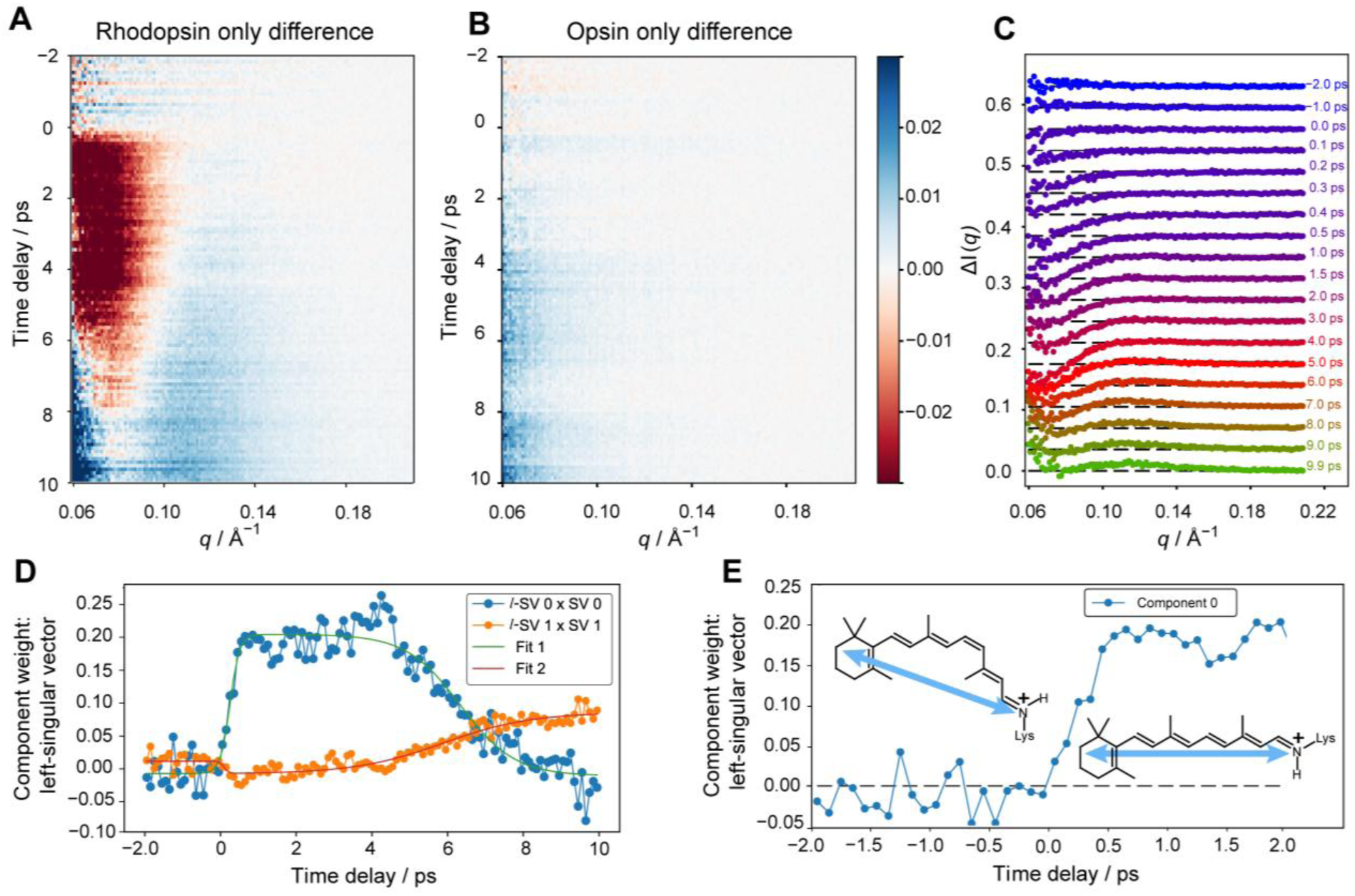
Femtosecond time-resolved X-ray scattering analysis of light-induced rhodopsin structural changes. **(A)** Experimental TR-XSS difference profiles (light–dark; *λ*=527 nm) for rhodopsin (10 mg mL^−1^ in 30 mM CHAPS detergent) at pH 7.4 and ambient temperature. The TR-XSS profiles are shown as a stack of 1D profiles, colored as a heatmap. Each horizontal line corresponds to a single scattering profile at a particular time delay (blue, positive; red, negative). **(B)** Corresponding data for retinal-free opsin apoprotein; the color-bar shows difference-scattering intensity in units of photons per pixel. **(C)** Examples of rhodopsin-minus-opsin double-difference profiles for representative times from –2 to +10 ps**. (D)** Singular-value decomposition (SVD) of double-difference profiles reveals two significant states. Amplitudes of the first two left-singular vectors (*l*-SV) versus time are fitted with a sum of exponentials. **(E)** Closeup for dominant left-singular vector during first two ps shows the most significant state rises within the first time point coinciding with 11-*cis* to all-*trans* isomerization of retinal.

### Resonant photon absorption as revealed by ligand-free opsin apoprotein

Crucially, the difference signal for rhodopsin at the low *q*-values (Fig. 2A) was absent for the control opsin apoprotein, which lacks the retinal cofactor (Fig. 2B). Besides light absorption by the retinal chromophore, nonresonant absorption by amino acid residues such as tryptophan is possible, e.g., as seen with bacteriorhodopsin ^23^. To address the question of resonant versus non-resonant absorption, we prepared retinal-free opsin which is not photochemically active. Contributions from absorption by the protein and detergent— i.e., not arising from retinal photoabsorption—were measured by collecting an additional dataset for the opsin apoprotein (Fig. 2B). The opsin difference profiles showed little to no scattering signal across the entire *q*-range, in striking contrast to the rhodopsin holoprotein. Double-difference scattering profiles were calculated to remove any small residual signals from nonresonant effects. The TR-XSS data for the opsin control prove that the ultrafast changes in the TR-XSS signal originate from resonant photon absorption by the retinal cofactor. Such control studies have rarely been conducted with an XFEL due to the limited beam-time availability.

Perhaps most surprising, our results imply the light-induced TR-XSS differences for rhodopsin are due to ultrafast (sub-picosecond) *cis–trans* isomerization of the retinal cofactor. Structural detection of retinal isomerization in real time has never been achieved for rhodopsin to our knowledge. The crucial opsin control (Fig. 2B) proves that energy transfer from retinal spreads throughout the rhodopsin holoprotein. The photonic energy (∼54 kcal mol^−1^) internally converts the protein to an energetically favorable state (photorhodopsin; see below) that transitions downhill in the multiscale reaction mechanism (Fig. 1B). Experimentally, increasing the pump (OPA) power density deposits excess energy to the surroundings ^24^, while maintaining the biological reaction pathway ^25^. From a biophysics viewpoint, the changes seen in the TR-XSS experiment owe to rapid *cis* to *trans* isomerization of retinal, causing the protein to evolve on its energy landscape, accompanied by greater internal energy of the photointermediates versus the dark state (Fig. 1C).

### Time-ordered reaction sequence in rhodopsin X-ray scattering

To better understand how the photointermediates yield differences in X-ray scattering, we carried out singular-value decomposition (SVD) of the experimental TR-XSS profiles (Fig. 2D, E). These profiles were obtained from class averages of scattering images based on their median intensity in a specified *q*-range (Fig. 3A). Initially, several hundred images were grouped into 100-fs time-delay bins (Fig. 3B, C). Based on shot intensities (Fig. 3A), we calculated the opsin minus rhodopsin double-difference profiles (Fig. 3D, E). The SVD results—the right singular vectors in the structural *q*-space and left singular vectors in the time domain—showed the scattering difference was dominated by two components (Fig. 3F). The largest component begins to rise within the first time points of the TR-XSS data (∼30 fs or ∼100 fs bins) (Figs. 2D, 3G), which overlaps formation of the primary photoproduct (∼90 fs) passing through the CI ^2^. This state grows until a plateau is reached at ∼500 fs, which persists until ∼4 ps after the optical pulse. Thereafter, the dominant component gives way to a second component, rising continually through the last time delay at 10 ps, at both 100-fs (Fig. 2D) and 30-fs binning (Fig. 3G). The conservative 100-fs binning ensured no information was lost by class-averaging the scattering profiles. The similar left singular vectors binned at 30 fs corroborated the results, despite higher noise levels, approaching the time resolution of ultrafast transient-grating studies (Fig. 3H) ^8^.

**Fig 3.**
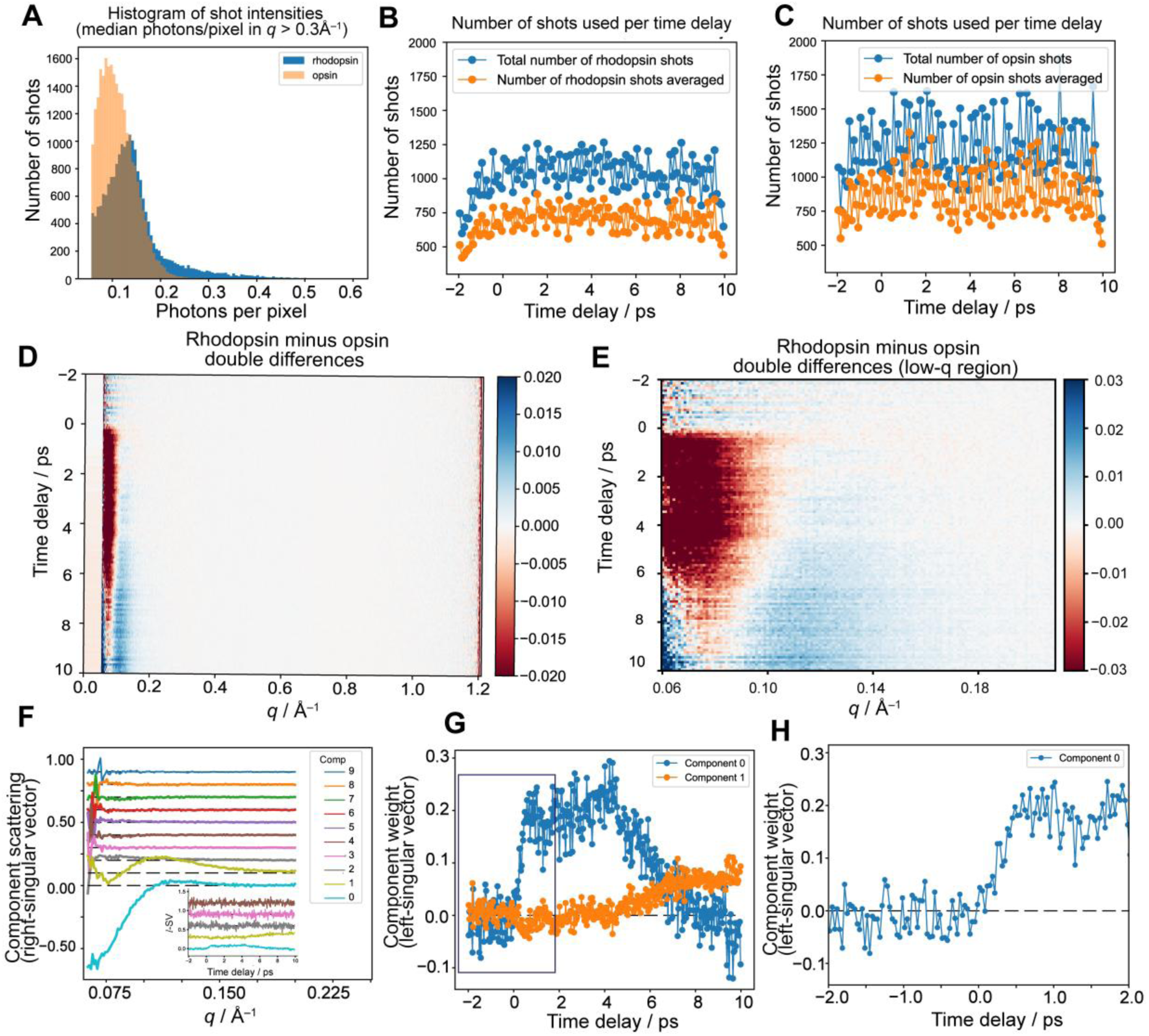
Time-resolved X-ray scattering analysis of ultrafast structural changes of rhodopsin compared to ligand-free opsin. **(A)** Histogram showing the number of scattering patterns versus the intensity per shot (photons/pixel) in the scattering vector range *q* > 0.3 Å^−1^ for opsin and rhodopsin. **(B, C)** Number of shots available for rhodopsin and opsin apoprotein experiments (blue) corresponding to each time delay (ps) with 100-fs binning plotted alongside number of shots actually averaged (orange) for analysis. **(D)** Heatmap shows rhodopsin-minus-opsin double-difference profiles at 100-fs binning. **(E)** Expansion of panel D between the scattering vectors *q* = 0.06 Å^−1^ and 0.22 Å^−1^. **(F)** The SVD components obtained when scattering profiles are binned at 30 fs. The first ten right-singular vectors (*r*-SVs) show only two components with significant structure. *Inset:* first five left-singular vectors (*l*-SVs) versus time delay, showing only two vectors with scattering features. **(G)** Expansion of data for first two left-singular vectors (*l*-SVs) from –2 to +10 ps. **(H)** Expansion of data for first *l*-SV is shown for –2 to +2 ps time delays.

Even though SVD vectors cannot always be assigned to distinct structural states, based on the timescale, the first component coincides with bathorhodopsin as the first stable photointermediate (<200 fs) (Figs. 2E, 3H). However, transient-grating experiments suggest the primary photoproduct (photorhodopsin) is formed beginning at ∼30 fs, whereas the average excited state lifetime is ∼90 fs for the population ^2^. For TR-XSS the longer time to complete the initial change (<500 fs) is likely because it probes the entire protein and not just the retinal binding pocket (see below). Surprisingly, the first SVD component decays concurrently with appearance of a second component (∼4 ps) within the optical lifetime for bathorhodopsin (tens of ns) ^6^. We attribute this transition to relaxation of the initial, impulsive structural distortions due to energy dissipation from the active site to the protein surface. Hence the second SVD component may represent a thermally relaxed state formed by the stabilization of the protein following the decay of the primary photoproduct ^6,8^. The remainder of the absorbed light energy must be internally redistributed within the protein to account for the time-ordered sequence of photointermediates in visual signaling (Fig. 1C) ^6^.

### Seeing rhodopsin photochemistry in real time with X-ray scattering

Accordingly, TR-XSS based on an XFEL (Fig 1A) provides a unique structural window into the ultrafast photochemistry central to dim-light vision. Optical spectroscopy (3, 7) and quantum-mechanical calculations (1, 6, 29) have established that the high quantum yield (0.67) of rhodopsin arises from a conical intersection (CI)—a shortcut of the electronic energy between the excited rhodopsin (11-*cis*) and ground photorhodopsin (all-*trans*) states (cf. Fig. 1C). However, a fundamental gap remains in translating this electronic phenomenon into structural reality. For rhodopsin after photon absorption, the CI point enables a barrierless (non-radiative) *cis*-to-*trans* isomerization of the retinal cofactor. The nuclear and electronic motions are not separable (due to vibronic coupling—failure of the Born-Oppenheimer, BO approximation) leading to the ultrafast reaction speed (Fig. 1C). Theoretically, the CI point involves a non-adiabatic singularity where electronic and nuclear motions become inextricably coupled during photoisomerization (nonadiabatic process). The nuclei are viewed as abruptly hopping from the excited to the ground-state potential (surface crossing). By contrast, for non-ultrafast processes the nuclear positions evolve gradually on a single potential surface (adiabatic theorem), so that adiabatic or BO states have an avoided crossing. Hence, the isomerization entails diabatic states and the quantum yield has been calculated using the Landau-Zener formula^26^.

Notably, the CI point predicts that electronic energy should be instantly converted into nuclear motion. Therefore, the specific structural manifestation of this conversion has eluded direct observation. Here we show that TR-XSS uniquely bridges this divide between quantum dynamics and structural biology. While X-ray scattering does not map the electronic potential energy surfaces directly, it provides a time-resolved sampling of the resulting nuclear coordinates. Atomic motions of retinal constitute the reaction coordinates of C11=C12 bond isomerization, including localized ethylenic stretching, hydrogen-out-of-plane (HOOP) wagging, and torsional modes (*7, 18*). The TR-XSS signal at early times are due to coherent nuclear motions of retinal coupled to the protein, which not only survive passage through the CI, but also drive its *cis*-*trans* isomerization (*2, 3, 6, 7*). Upon vertical photoexcitation (Franck-Condon states), the vibrations on either side of the CI lead to molecules passing to the photoproduct side (all*-trans*) (quantum yield = 0.67), while the remainder (0.33) simultaneously populates the reactant side (11-*cis*) (called recovery of ground-state bleach, RGB). Importantly, the RGB is a signature of vibrational coherence—it implies passage through the CI point (*7*). It is caused by photoexcited but unreactive retinal molecules which undergo internal conversion to the vibrationally hot 11-*cis* ground state (*7*). Consistent with this proposal, the first singular vector rises with a time constant (127 fs) closely resembling the RGB value (∼100 fs) in the kinetic analysis (*7*). It likely includes photorhodopsin coupled to the reactive surface-crossing, together with bathorhodopsin. Our interpretation is consistent with the second left-singular vector (appearing at ∼4 ps) having a rising time constant of 1.2 ps similar to thermally relaxed bathorhodopsin (∼2.1 ps) (*7*).

In our experiment, we come close to the rarely observed theoretical CI point and measure the aftereffect of the potential surface crossing on the protein dynamics (*7, 19*). The scattering components suggest transfer of delocalized vibrations within the protein manifesting as a coherent, ballistic rearrangement of the protein scaffold that emerges immediately (<500 fs) from the non-adiabatic transition (Fig 3H). This observation confirms that the retinal electronic relaxation is not dissipated thermally but is instead efficiently channeled into a directed mechanical trajectory, subsequently coupling with solvent on picosecond timescales. Our TR-XSS measurements complement serial crystallography (TR-SFX) (*20*) while avoiding crystal-lattice forces, e.g., as seen for photoactive yellow protein (PYP) (*21*). In TR-SFX the isomerization is fully complete before the first time point (at 1 ps) (*20*), corresponding to the earlier cryotrapped bathorhodopsin structure (*33*). However, our TR-XSS study has many time points (120 compared to 3 for TR-SFX) and we resolve the *cis–trans* isomerization of retinal as it occurs in real time (Figs. 2E, 3H)—the defining moment of visual photochemistry.

### Molecular simulations elucidate ultrafast X-ray scattering of visual receptors

We conducted all-atom molecular simulations of rhodopsin equilibrated in detergent micelles with an explicit solvent ^27^, to structurally interpret the ultrafast changes ^4,7^. Our MD approach emulates more exact quantum-mechanics/molecular-mechanics methods corresponding to adiabatic states of the ground-potential energy surface (beyond the CI point or avoided crossing) (Fig. 1C) ^2,20,28^. Although pushing the quantum limit ^1,20^, it more fully captures the structural diversity inherent to the ensemble of rhodopsin molecules ^27,29^. We conducted 18 independent, ∼600-ns simulations of dark-state rhodopsin in CHAPS detergent micelles for an ensemble of 10,000 starting configurations . From each configuration, we performed two paired simulations—one where we continued the simulation directly from the saved state, and a second where we classically modeled the photon absorption. The *trans* state of the retinal was favored by transiently imposing a potential barrier (60 kcal/mol) to destabilize the *cis* state. Thereafter energy was added to the isomerized state to approximate the adiabatic energy gap versus the dark state . To separate intrinsic protein dynamics from retinal isomerization–induced conformational changes, we calculated MD-derived difference electron density maps Each atom in the separate trajectories was represented by a Gaussian function whose amplitude was determined by its corresponding number of electrons to generate electron density maps from the paired simulations ^30^. Difference electron density maps (light–dark) were generated with a 100-fs time resolution and were averaged over all the trajectories ^30^. The resultant single final map for each time point was superimposed on the dark-state rhodopsin coordinates (Fig. 4A). The large number of time points (Fig. 4B, C), revealed the ultrafast displacements of conserved amino acids coupled to the retinal isomerization. Importantly, we found the largest initial electron density difference was located close to the C11=C12 double bond and the retinylidene protonated Schiff base (PSB) of Lys^296^ (Fig. 4A), migrating toward the crucial β-ionone ring by 200 fs. The sum of difference electron densities near the active site (Fig. 4C center) shows that local structural changes grow until ∼500 fs, after which they are completed, similar to the experimental TR-XSS data.

**Fig 4.**
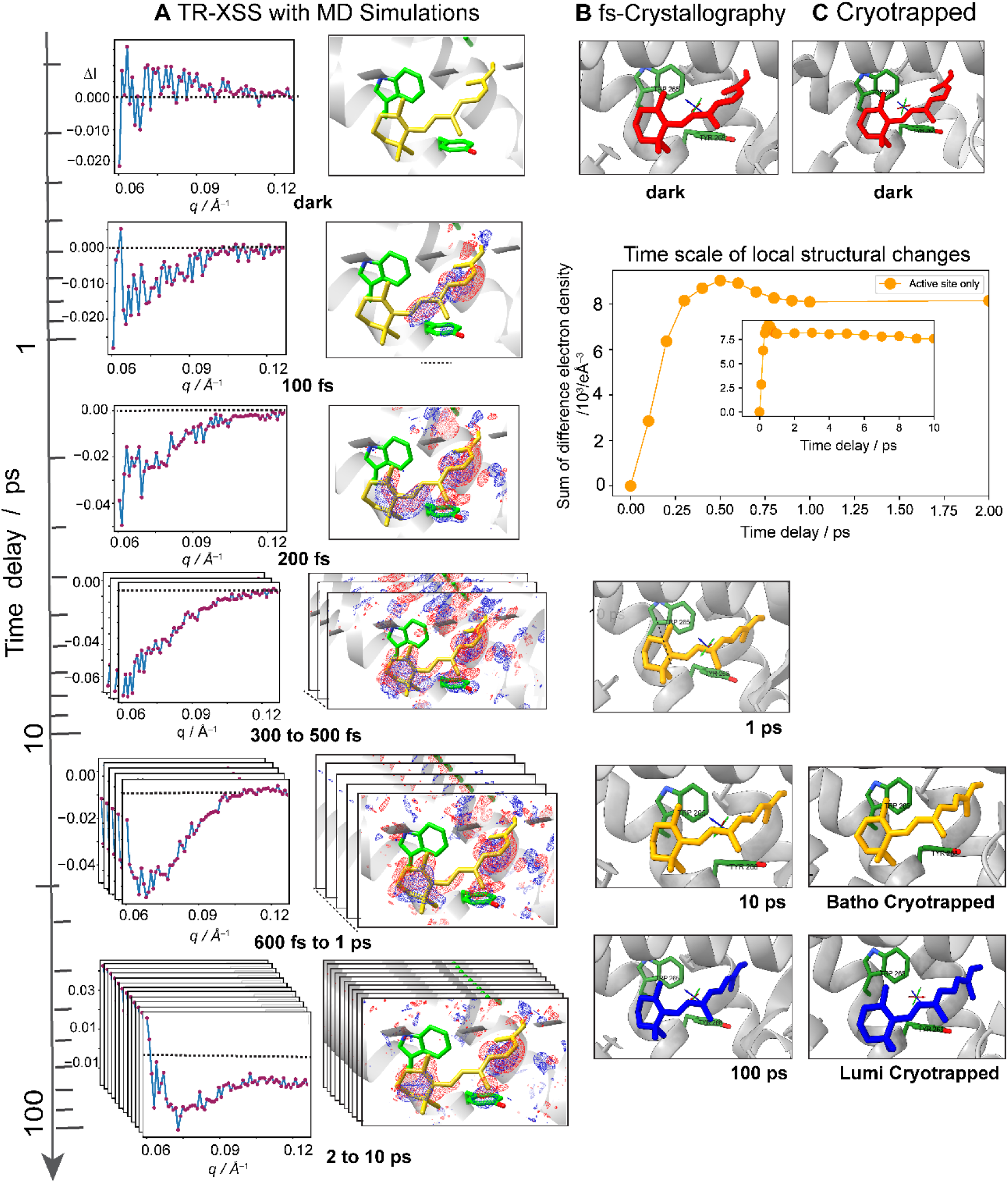
Timeline of X-ray scattering analysis of rhodopsin structural changes compared to photointermediates. **(A)** Light-minus-dark scattering profiles (left panels) showing evolution of averaged difference electron densities (right panels) from ∼8000 paired MD simulations superimposed on retinal, Tyr^268^, and Trp^265^ of dark-state rhodopsin from a random single trajectory. Positive difference density is shown by blue-colored mesh and negative by the red mesh around Tyr^268^ and Trp^265^ residues. The retinal isomerization was revealed by negative density at the 11-*cis* bond and positive density for the *trans* conformation. Density maps are contoured at 40 σ. By 500 fs evolution of the difference density around the Tyr^268^ residue is completed. **(B)** Complementary view of structures from time-resolved serial femtosecond crystallography. PDB codes: 7ZBE (dark), 8A6C (1 ps), 8A6D (10 ps), and 8A6E (100 ps) **^15^**. **(C)** X-ray structures of cryotrapped photointermediates. PDB codes: 1U19 (dark), 2G87 (bathorhodopsin, ps–ns scale), and 2HPY (lumirhodopsin, µs–scale) ^14^. *Center*: sum of difference electron densities within 5 Å of retinal versus time delay in ps, showing stabilization at ∼500 fs. *Center inset:* expansion of sum for entire 10-ps range. Note the TR-XSS experiments have greater time resolution than crystallography. Figure prepared with CHIMERA ^52^.

Next, to enable fitting of atomic models to the experimental TR-XSS data, we customized a module of DENSS ^31^ for implementation with the difference profiles (unlike conventional routines). Direct calculation from explicit coordinates proved highly sensitive to background mismatches between the equilibrated simulation box and the vacuum-cooled liquid jet. To isolate the biological signal, we utilized an implicit solvent model (DENSS) that treats the solvation environment as a statistical continuum. Robust fitting required allowing the implicit solvent density parameters—specifically excluded volume density *c*_1_ and hydration shell contrast *c*_2_ —to vary. These parameters function as mean-field proxies, absorbing ensemble-averaged protein-solvent cross-terms and experimental baseline offsets. This approach circumvents the systematic errors of explicit calculations, confirming the protein structural trajectory is consistent with the experimental data (Fig. 5A).

**Fig 5.**
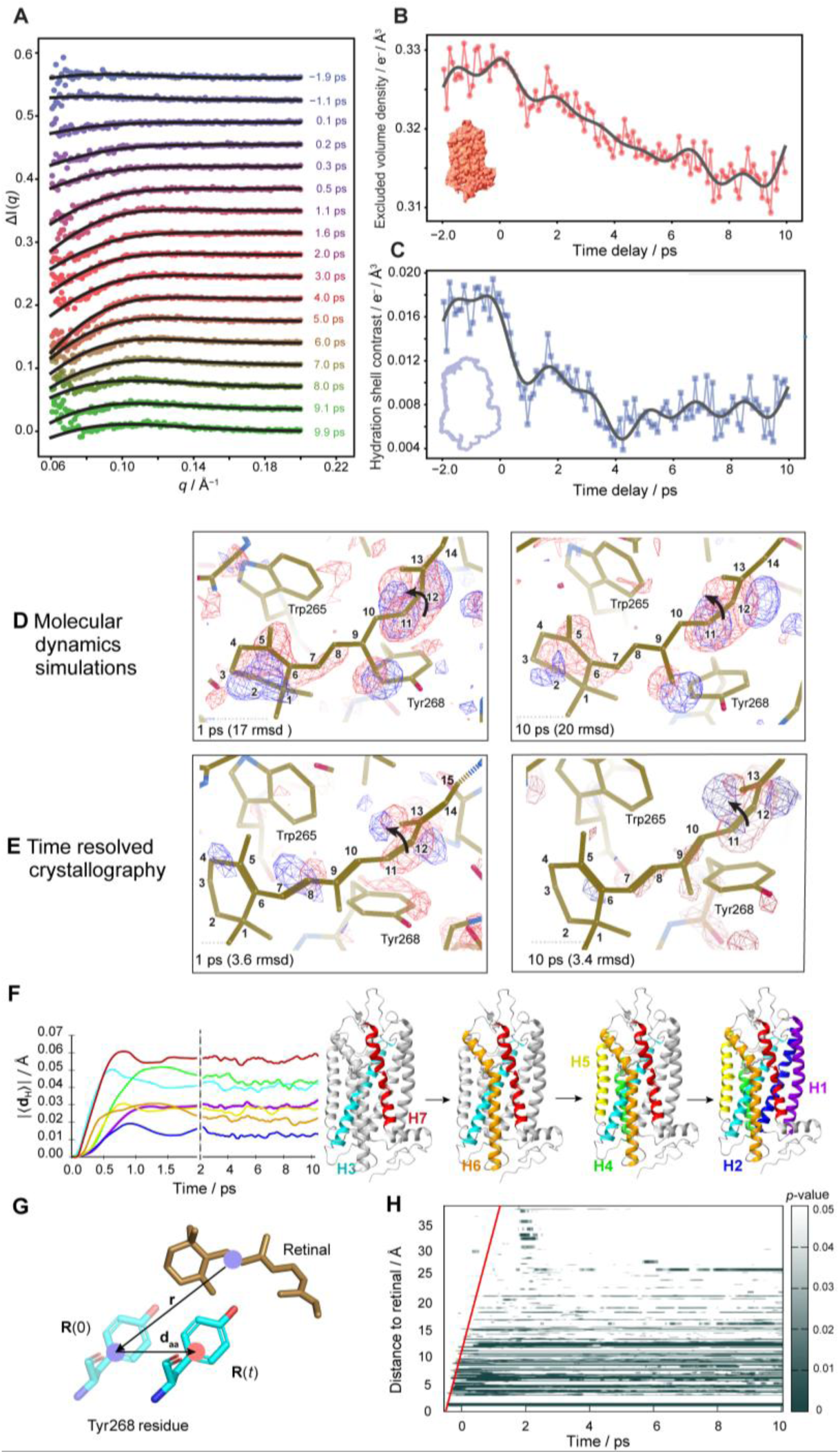
Integrated DENSS fitting and MD simulations elucidate ultrafast rhodopsin dynamics. **(A)** Rhodopsin minus opsin experimental scattering profiles are fit to the calculated difference-scattering profiles (black) obtained from dark and light-activated coordinates for representative time points from a single trajectory (out of 8000 MD trajectories) using DENSS ^31^. **(B and C)** Time sequence of optimized fitting parameters (in e/Å^3^) obtained from DENSS fits of rhodopsin light activate scattering profiles, namely, excluded volume density (bulk solvent) (panel B), and hydration shell contrast (panel C). For visualization purposes, a low-pass filter was applied to the data (solid line). *Left insets*: schematics of excluded volume (panel B, red) and hydration shell (panel C, blue). The red surface is made using CHIMERAX ^53^.**(D)** Averaged difference (dark–light) electron densities from MD simulations overlaid on dark retinal structure (red, negative; blue, positive difference electron density). *Top left:* difference at 1 ps and *bottom left:* 10-ps timescales. **(E)** Difference electron densities overlaid on the retinal from inactive dark-state TR-SFX crystal structure (PDB code: 7ZBE). *Top right:* panel is calculated from 1-ps crystal structure (PDB code: 8A6C) and *bottom right:* 10-ps crystal structure (PDB code: 8A6D). Both the simulations and crystallography show negative to positive electron density differences due to the C11=C12 bond isomerization. The MD simulations show greater motion of the crucial β-ionone ring. Figure prepared with Coot ^54^. **(F)** *Left:* time-dependent average displacement of center of mass |〈**d_H_**〉| of helices versus time after retinal isomerization. *Right:* sequence of transmembrane helical motions; H1, purple; H2, blue; H3, cyan; H4, green; H5, yellow; H6, orange; and H7, red. **(G)** Characterization of motions of amino acid residues as projection of center-of-mass displacement **d_aa_**(*t*) onto retinal distance vector r at time zero. Here R(0) and R(*t*) denote center of mass of the amino acid residue at times 0 and *t*, respectively. **(H)** Map of *p*-values for each of the residues sorted by their distance to retinal plotted against time. The *p*-values indicate statistically significant motion calculated for COM displacements for light versus dark simulations.

Physically the solvent environment remains effectively static on this sub-picosecond timescale. However, the retinal isomerization triggers immediate, ballistic nuclear motions within the binding pocket (Fig 4A). Our implicit solvent analysis captures the modulation of the scattering interference pattern against the fixed solvent background by these localized atomic displacements. These optimized parameters (Figs. 5B, C) begin to change within <100 fs, mirroring the first SVD component (Fig. 2A). This ultrafast timescale precludes thermal expansion, as heat dissipation from the buried chromophore to the bulk solvent requires picoseconds ^32^. Given the negligible opsin TR-XSS signal (Fig 2B), the solvent parameter changes initiated by the retinal isomerization, are driven by the protein-solvent scattering cross-terms ^33^. Here the varying density parameters are mathematical proxies for these changing cross-terms, effectively mapping the interaction between the impulsive protein kinetics and its static solvent cage. We leveraged this interaction to overcome the detection limits of wide-angle scattering (WAXS), where minute signals are often obscured by destructive interference and solvent noise. By contrast, our low-*q* (SAXS) measurements benefit from intrinsic signal amplification: the protein-solvent cross-terms sum coherently, generating constructive interference. This magnification allowed unprecedented detection of ultrafast nuclear motions via the hydration shell proxy, unlike previous XFEL studies relying primarily on WAXS ^34–37^, revealing structural dynamics that remain indiscernible in the wide-angle regime.

Compared with time-resolved crystallography ^15^, greater atomic displacements occur within the first 10 ps due to absence of crystal-lattice forces (Fig. 4B, C, 5D, E). Our simulations indicate sub-picosecond motions of key residues near the retinal binding pocket — Tyr^2^^68^, Trp^2^^65^, Ala^1^^17^ and the Cys^1^^10^–Cys^187^ disulfide bond that serve as precursors to later intermediates ^38,39^. Within the first 500 fs after photoisomerization, the retinylidene C9-methyl group interacts with the Tyr^2^^68^ phenolic ring (Fig. 5D) of a key functional hydrogen-bonding network ^40^. Both the retinal C9-methyl group and the Tyr^268^ residue are crucial to the catalytic activation of transducin by light-activated rhodopsin (MII state) ^40,41^. Hence we propose that ultrafast steric interaction of the Tyr^268^ phenolic ring with the C9-methyl group initiates rhodopsin activation in the signaling cascade.

These ultrafast changes are absent in the cryotrapped bathorhodopsin and lumirhodopsin structures at the longer ns-timescales (Fig. 4C) ^14,42^. The transmembrane helices most perturbed by retinal isomerization moved within hundreds of femtoseconds, peaking around 500 fs and 1.2 ps depending on the helix (Fig. 5F). Significant global protein movements were characterized by sorting the amino acid residues by their distance to retinal (Fig. 5H) and explained by their center-of-mass displacement **d_aa_**(*t*) from retinal (COMD) versus time after light excitation (Figs. 5G, H). We found the perturbation due to retinal isomerization spreads at a rate of ∼21 Å·ps^−1^, approximately the speed of sound within the protein matrix ^43,44^. The nonrandom helical motions indicate they are controlled by the protein structure. For GPCRs, conserved sequence motifs create weak spots in transmembrane helices involving helical motions, which require elimination or stabilization through mutation before crystal structures can be determined ^45,46^. Our analysis effectively links the electronic energy of the conical intersection to the mechanical motions that initiate the earliest steps of the visual signaling cascade.

### Comparing rhodopsin to other photoactive proteins

It is significant that rhodopsin differs from other non-light-driven proteins such as myoglobin and hemoglobin that have been investigated using TR-XSS methods ^37,44,47^. For rhodopsin the system transitions into an excited electronic state yielding *cis–trans* isomerization of the ligand ^2,3^. This behavior contrasts with myoglobin and hemoglobin, which biologically do not involve light-driven reactions. In those cases, light is used artificially to drive the dissociation of CO from the heme cofactor, followed by its re-binding ^43,48^. Yet the energy landscape with hills and valleys ^49^ is populated without any additional input of energy yielding the active state. By contrast, the rhodopsin photointermediates entail retinal isomerization, protein conformational changes, and solvent reorganization by energy coupling to the surroundings. Nuclear motions of retinal are captured when passing through the transition state of the photoproduct. Advances in next-generation X-ray sources ^50^ will motivate applications to proteins with caged photoactive ligands, membrane lipids, and optogenetics ^11,51^. Remaining questions include how X-ray scattering detects both nuclear and electronic motions and how internal energy of rhodopsin is partitioned in the visual process. Future insights will doubtlessly emerge at the nexus of ultrafast dynamics with photochemistry and structural biology in understanding biological mechanisms.

## Methods

### Rhodopsin purification and sample preparation

Retinal disk membranes (RDM) were isolated from bovine retinas as described ^55^. Unless otherwise stated all procedures were done at 4 °C under dim red light or darkness. Electronic (UV–visible) absorption spectra for both the dark and light-exposed states of rhodopsin were acquired at room temperature with a modified Cary-50 UV-visible (single-beam) spectrometer (Varian, Inc.; Palo Alto, CA). Purified bovine RDM typically had unregenerated *A*280/*A*500 absorption ratios < 2.9. The RDM samples were further characterized by their regenerability with 11-*cis*-retinal or 9-*cis*-retinal ^55^ with typical results of 5 % bleached, 97% regenerable, and regenerated *A*280/*A*500 spectral ratio ∼2.0. To produce the required large amounts of solubilized rhodopsin (∼1000 mg per beam time) a new detergent protocol was developed. Choice of the detergent entailed increasing the protein molar concentration relative to the pump laser fluence while supporting the photochemical functionality of rhodopsin ^55^. The zwitterionic detergent 3-[(3-cholamidopropyl)dimethylammonio]-1-propane sulfonate (CHAPS) (Anatrace; Maumee, OH) was used owing to the small micelle aggregation number (∼10 versus ∼120 for *n*-dodecyl-β-D-maltoside) and high critical micelle concentration (cmc, 8 mM), yielding a thin belt around the protein hydrophobic domain. The rhodopsin in RDMs was first solubilized in the cationic detergent dodecyltrimethylammonium bromide (DTAB) (Sigma-Aldrich; St. Louis, MO) and purified by hydroxyapatite (HA) column chromatography ^55^. After HA column purification the final *A*280/*A*500 ratio of < 1.9 indicated a highly pure rhodopsin sample. The detergent was then exchanged from DTAB to CHAPS by size-exclusion chromatography (SEC). The rhodopsin/DTAB solution was applied to a Sephadex G50 column and eluted with the CHAPS detergent buffer (typically 30 mM CHAPS, 15 mM phosphate, pH 6.7). Characterization of the CHAPS-isolated rhodopsin entailed dynamic light scattering; together with SDS gel-electrophoresis and native mass spectrometry. The high-purity (*A*280/*A*500 <1.9) fractions were then pooled and concentrated using 30-kDa centrifugal filters. The final rhodopsin/detergent molar ratio of the sample was nominally ∼1:40 and the final concentration used at the beamtime was typically 10 mg mL^−1^, yielding a homogenous preparation. The samples were flash-frozen in liquid nitrogen in 15-mL portions and stored at –80 °C until thawed just prior to use. Typically, 800–1200 mg of the rhodopsin/CHAPS samples were produced for each beamtime.

### Time-resolved X-ray solution scattering data acquisition

The diffract-then-destroy principle ^56^ was used to acquire the time-resolved X-ray solution scattering (TR-XSS) data with a pump-probe scheme. The reported studies used bovine rhodopsin reconstituted in CHAPS detergent (typically 10 mg mL^−1^) and the apoprotein opsin under identical conditions. Our experiments were conducted using an X-ray free-electron laser (XFEL) with self-amplified spontaneous emission (SASE) at the Coherent X-ray Imaging (CXI) beamline of Linac Coherent Light Source (LCLS) at SLAC National Accelerator Laboratory during beam time LU18. Additional supporting studies were conducted at LCLS during beam times LE71, LM59, LN60, LP99, and LW75. For LU18 the XFEL pulse duration was 40 fs at 9.5-keV energy (*λ* = 1.3 Å), with a 120-Hz repetition rate in the microfocus instrument in its standard configuration at CXI. The TR-XSS data were acquired using a Cornell-SLAC pixel-array (CSPAD) detector. The detector was moved back to 0.507 m which allowed us to collect the *q*-range between 0.06 Å^−1^ and 1.2 Å^−1^ (*q* = 4π sin *θ*/λ, where the scattering angle is 2*θ* and *λ* is the X-ray wavelength). The samples were delivered to the X-ray interaction region using a gas dynamic virtual nozzle (GDVN) or 3D-printed nozzle (50-µm inner diameter) ^57^. The average liquid jet-diameter varied between 3 µm and 5 µm. The flow rate of the rhodopsin or opsin/CHAPS detergent solution varied between 25 and 35 µL min^−1^. Samples were continuously replaced (jet velocity up to 70 m s^−1^) with the pump-laser system triggered at 60 Hz, interleaving laser-on and laser-off measurements. To initiate the rhodopsin photochemistry, for the actinic light source (pump) we used an optical parametric amplifier (OPA) driven by a Ti:Sapphire laser (TOPAS; Spectra-Physics; Santa Clara, CA) with a ∼70-fs pulse width at 527 nm in a collinear geometry. The laser was focused having a beam diameter of 0.195 mm (FWHM) with fluence of 1.33 mJ mm^−2^ at the interaction point. The OPA generated pulses of 30 uJ ± 1% pulse energy, which were attenuated with a polarizer and a half-wave plate. Photoselection of rhodopsin molecules with the retinal transition dipole moment aligned along the electric field of the actinic light pump was minimized by using circularly polarized pump excitation. The scattering images were collected with –2 to +10-ps nominal time delays between the optical pump and X-ray probe, in intervals of 100 fs. Each interval had an average of ∼1250 shots. The profiles were also binned and grouped at 30-fs time delays, which did not affect the conclusions (Fig. 3 of main text). The light–dark difference profiles spanned *q*-values from 0.06 to 1.2 Å^−1^. On-the-fly data reduction and analysis was done with a customized version of OnDA ^58^ developed for pump-probe TR-XSS studies. We used OnDA to monitor the time-resolved difference scattering profiles directly from shared memory as the data were collected, before being written to disk. This capability informed our experimental strategy during the beam time by allowing us to optimize the alignment of the liquid jet and the spatial overlap of the pump laser.

### Analysis of light-minus-dark X-ray difference scattering profiles

After radially averaging the profiles from each experimental run, the scattering profiles were binned corresponding to the time delay for both the holoprotein rhodopsin and the retinal-free apoprotein opsin. For both the rhodopsin and opsin proteins, the pump-probe time delays were obtained using the time tool ^22^ at the CXI beamline of LCLS. For a given sample, the light-minus-dark difference profiles were obtained showing the changes in the radial scattering before and after the pump laser event. The dark profiles were obtained by radially averaging the images acquired when the pump laser arrived after the X-ray pulse. The dark scattering profiles were first binned according to the median intensity in the high *q*-range between 0.6 and 1 Å^-1^. This region was selected because it had the least variation among all the shots. After binning the data, the outliers in each bin were filtered out before averaging. The outliers with more than two standard deviations from the median scattering profile were rejected based on the root mean square of the summed residuals as *R*_sqr_ = { ∑*_q_*[*I_i_*(*q*, *t*_0_) − ⟨*I*(*q*)⟩]^2^}^1/2^, where *I_i_*(*q*, *t*_0_) is the *i*th scattering profile and ⟨*I*(*q*)⟩ is the median scattering profile for the given intensity bin. In the experiment, the scattering signal is a function of the liquid-jet thickness, sheath gas from the GDVN, and incident X-ray fluence. The X-ray fluence and jet thickness varied stochastically whereas the sheath gas remained essentially stable. Because the jet thickness was unknown from shot-to-shot, and the relative contributions from the liquid and the gas varied, we grouped (class averaged) the patterns according to the overall scattering signals before taking differences. The scattering intensity in the aforementioned *q*-range was used as an indicator for the product of the incident X-ray intensity and jet thickness. The light-minus-dark scattering profile differences were acquired between the shots with identical experimental conditions, such as liquid jet thickness and X-ray intensity, which used an alternating light/dark pulse scheme with the 120-Hz XFEL, such that pumped images and dark images were from identical experimental conditions. We matched the pumped radial-scattering profiles to the dark profiles belonging to the same median X-ray intensity bin. To account for variation within an intensity bin having several scattering profiles, each average dark-profile was then fit to the pumped profile before being subtracted. The above scheme of light-minus-dark subtraction was followed for the radial scattering profiles of both rhodopsin and its apoprotein opsin (the ligand-bound and apo forms).

### Rhodopsin-minus-opsin double-difference scattering profiles

To compare rhodopsin to its apoprotein at positive time delay after the laser pump, we subtracted the above light-minus-dark difference scattering profiles of the apoprotein from that of rhodopsin. Profiles were binned according to the median intensity of the pumped radial scattering profiles in the high *q*-range (between 0.3 and 1 Å^−1^). The *q*-range was chosen since both the ligand-bound and apo forms of the protein (the apoprotein opsin and rhodopsin holoprotein) had similar scattering intensities in this range. After binning, the outliers in each bin were filtered before averaging. The averaged negative time-delay difference profiles were subtracted from the positive time-delay difference scattering profiles for opsin and rhodopsin respectively to remove small artifacts in the signal, as the negative-time difference scattering profiles should exhibit a negligible signal. A histogram of scattering intensities (photons per radial pixel) (Fig. 3A of main text) showed that the majority of scattering images for both opsin and rhodopsin had intensities less than 0.2 X-ray photons per radial pixel. The corresponding opsin and rhodopsin difference profiles with the same shot intensities (< 0.2 photons per radial pixel) were matched and the opsin difference profiles were fit to the rhodopsin difference profiles using a least-squares routine. The fitted opsin difference radial scattering profiles (Fig. 2B of main text) were subtracted from rhodopsin difference scattering profiles (Fig. 2A of main text) to obtain the double-difference scattering profiles of rhodopsin minus opsin (Fig. 3D, E of main text).

### Kinetic analysis of double-difference X-ray scattering

Analysis of the structural changes entailed singular value decomposition (SVD) of the double-difference rhodopsin-minus-opsin scattering profiles as described above. The SVD of the scattering profiles produced left-singular vectors (*l*-SVs) (Fig. S5A), right-singular vectors (*r*-SVs), and singular values. The *r-*SVs had the same structure as the difference scattering profiles and the *l*-SVs gave the time dependence of the *r*-SVs. Each of the singular values gave the relative weights of the singular vectors contributing to the double-difference rhodopsin-minus-opsin profiles. The SVD analysis revealed two major *r*-SVs as evidenced by the magnitude of the singular values (Fig. S5C). The two most prominent *l*-SVs weighted by the singular values gave the temporal behavior of the *r*-SVs (see Figs. 2D, E and 3G, H of main text). The time sequence of the *l*-SVs was used to relate the energy propagation by global structural perturbations to the lifetimes of optically detected photointermediates (see main text). The growth and decay of the two *l*-SVs were fit to a sum of two logistic functions, one for the rise and another for the *l*-SV decay. The general form of the logistic function is: *y* = *a* / [1 + *e*^−*b*(*t*−*c*)^]. In this expression, *a* is the maximum value at the plateau of the rise of the *l*-SV, where *b* is the steepness of the rise or decay of the *l*-SV, and *c* defines the inflexion or midpoint of the rise or the decay of the *r*-SV. The time-constants of the rise/decay of the *l*-SVs were defined as 1/*b*, which were obtained from the fits of the sum of logistic functions to the two prominent *l*-SVs. This model-free approach is very similar to the fits of the time-dependent singular vectors ^59^ using an exponential decay function convoluted with a Gaussian instrument response function ^37^. Our time-dependent *r*-SVs were better represented numerically by a logistic function as each of them have both a rise and a decay.

### Molecular dynamics simulations

The crystal structure of dark-state bovine rhodopsin (PDB code: 1U19) ^13^ was used as the starting structure for the simulations in CHAPS detergent micelles. Rhodopsin was inserted into a 109-Å cubic box with its geometric centroid at the origin and its calculated membrane normal as in the Orientation of Proteins in Membranes (OPM) database ^60^ aligned to the *z*-axis. The water boxes were large enough so that the spatial envelopes could segregate the solvent layer of the solute from bulk water ^43,61^. The initial conformations of the micellar detergents were randomly chosen from the CHARMM-GUI website library (http://www.charmm-gui.org/?doc=archive&lib=lipid) ^62^. In total, 40 CHAPS molecules were iteratively placed around the protein inside a cylindrical shell with height equal to the thickness of the receptor transmembrane region (21.1 Å) ^60^. The inner radius was equal to the maximum radius of the receptor in the TM region (22.9 Å), and the outer radius was equal to the inner radius plus the average detergent length in the library (25.7 Å). Detergent placements were accepted or rejected based on the minimum distance to other protein and detergent atoms (cutoff < 1.75 Å), and the relative orientation of the tail and headgroup versus the micelle center. The resulting protein/micelle complexes were solvated with 39,378 water molecules and 150 mM NaCl (plus one neutralizing sodium ion). This process was repeated to generate a total of 18 solvated micelle systems (simulations).

Before investigating the activation dynamics of the protein, each protein/micelle system had a production run of ∼600 ns in the dark state, for a combined simulation time of over 10 μs. The simulation time of 10 μs was required to capture the relaxation of structural diversity in the dark state of rhodopsin ^27^. The starting points of the (paired) activation simulations (described below) were obtained from this dark-state conformational ensemble. We conducted molecular dynamics (MD) simulations on the Summit computer at Oak Ridge National Laboratory (ORNL), using the NAMD simulation package (version 2.13) ^63^ with the CHARMM36 force field ^64–66^ and retinal parameters ^67^. The velocity Verlet integrator was used for dynamics propagation and all hydrogen-containing bonds were constrained with the SHAKE algorithm ^68^ to allow for a 2-fs timestep. Treatment of long-range electrostatics entailed the smooth particle-mesh Ewald (PME) summation method, employing a grid spacing of 1 Å. The real-space cutoff for nonbonded interactions was 12 Å. The dark-state rhodopsin/micelle systems were minimized and initially equilibrated in the canonical *NVT* ensemble with position restraints on the protein and the detergents. The detergent restraints were then switched off and the micelle equilibrated for 100 ns in the *NPT* ensemble ^61^. After that, the protein restraints were tapered off over 1 ns before entering the production runs. The *NPT* conditions (1.01 bar and 310.15 K) were maintained with a Langevin piston (oscillation period = 200 fs; damping timescale = 100 fs) and a Langevin thermostat (damping coefficient = 2 ps^−1^).

To explore rhodopsin activation at ultrafast timescales, we ran 10,080 paired (dark and light-activated) simulations. As the experimental timescales were short (fs–ps), the feasibility of running many activation trajectories improved the convergence of the results. For the paired simulation systems construction, the starting configurations were taken from the dark-state rhodopsin/micelle systems as described above. Briefly, atomic coordinates and velocities were extracted from the dark trajectories every 1 ns, for a total of 10,080 initial configurations. Each starting configuration was forked into a pair of (initially) identical simulations (dark and light-excited), beginning with identical coordinates and velocities, such that cancellation of signals not related to light excitation (e.g., water, lipids, remainder of the protein) was optimal when taking the paired difference between the two conditions. This noise reduction scheme was critical, as the signal of interest can be several orders of magnitude smaller than the noise. For each paired system, one simulation (dark) was left unchanged in the dark state. The other simulation (light) was light-excited classically by inducing retinal isomerization from 11-*cis* to all-*trans* and increasing the retinal kinetic energy. The original potential for this torsion is shown in pink. To induce retinal isomerization, we temporally modified the potential of the C10-C11=C12-C13 dihedral. and adjusted the force field parameters to impose a 60 kcal mol^−1^ energy barrier on the *cis* configuration of the C10–C11=C12–C13 torsion potential during the first 0.5 ps to favor the *trans* configuration (as shown in yellow) ^69^. The choice of torsional potential was validated against NMR results for the later photointermediates ^69^, and also agrees with the electronic excitation energies for So to S1 transition for rhodopsin calculated in previous QM/MM studies by Battista et. al ^70^. With the energy barrier, retinal isomerization was completed within ∼200 fs, as characterized experimentally ^5^. After 0.5 ps the original torsion potential was restored, and the simulation continued. At this point, we scaled up the velocities to capture energy released when the retinal would have relaxed to its ground state. This approach was sufficient for roughly 79% of the trajectories to undergo *cis*-*trans* transition, showing that we retained the effects of the protein environment. To increase the kinetic energy of retinal, the initial velocities of its polyene chain

atoms were rescaled as *vi*\*= [(*K*ret + *E*add) / *K*ret]^1/2^ *vi* where *vi*\* is the initial velocity of the *i*th atom, *vi* is its rescaled velocity, *K*ret is the sum of kinetic energies of the polyene atoms, and *E*add is the energy of one actinic photon as used in the experiment (∼54.25 kcal mol^−1^ at 527 nm). Notably, the addition of *E*add alone —without modifying the potential of the C11=C12 torsion— was not enough to induce retinal isomerization in this classic model. Each paired simulation was run for 10 ps, with the same simulation parameters used for the parent dark-state trajectories. Only the temperature control was different, as the thermostat was turned off to preserve the correct short-time dynamics. From the paired simulations, atomic coordinates were stored every 100 fs and analyzed as described below.

### Analysis of molecular dynamics trajectories

Subsequent trajectory processing and analysis was performed on BlueHive, the Linux cluster at the University of Rochester. For each of 10,080 paired (dark and light-activated) simulations., the individual frames were superimposed onto the dark-state crystal structure (PDB code: 1U19)^13^ using a Kabsch least-squares algorithm ^71,72^ with the transmembrane (TM) α-carbons as the alignment selection. This procedure minimized the deviation of TM α-carbons from the dark state structure thereby allowing comparison of the time-dependent structural changes within the ensemble of trajectories for a given time point. Torsions along the retinal polyene chain were scanned and the simulation pairs with deviations larger than 90° from the *trans* configuration (or *cis* configuration of the C10–C11=C12–C13 torsion in the dark trajectories) were excluded (20%). In total, 8,016 dark/light trajectory pairs were included in the subsequent analyses. All reported averages were computed by taking the difference between the paired dark and light frames as a function of time and calculating the mean over 8,016 differences. Error bars indicate the mean ± standard error (SE) computed by treating each difference as a single data point. Dark controls were calculated by randomly pairing 10,000 dark trajectories and taking their difference as a function of simulation time. Statistical significance was assessed through paired *t*-tests comparing the dark and light sets at a given time point, with a significance level of 0.01. All estimated *p*-values were corrected for multiple hypothesis testing using the Holm-Šídák algorithm implemented in StatsModels ^73^. Unless otherwise stated, all analyses were conducted using in-house code available upon request or existing analysis tools included in LOOS ^71,72^. The LOOS source code is available for download on GitHub (http://github.com/GrossfieldLab/loos). Trajectory visualization, image rendering, and data plotting were done with VMD (version 1.9.3) ^74^, PyMOL (version 2.1.0) ^75^, and Gnuplot (version 5.0), respectively.

We computed the time-resolved electron densities of the protein and the ligand using the EMAN2 program ^30^ where each atom in the separate trajectories (from the paired simulations) was represented by a Gaussian function whose amplitude was determined by its corresponding number of electrons. The PDB files were generated containing a subset of atoms from the aligned dark and light trajectories using the *subsetter* tool from LOOS ^71^. Next, the electron density map of each PDB file was computed with the *e2pdb2mrc* tool, while light–dark difference maps were generated with the *e2proc3d* tool ^30^. The resulting maps were averaged using the *e2buildstacks* tool and the *e2proc3d* tool ^30^ with the *--average* flag. The consecutive differences of averaged light-minus-dark differences were obtained using Chimera ^52^. Summing of light-minus-dark electron densities (Fig. 4 center of main text) and the consecutive differences (i.e., current time point minus previous time point) were done using DENSS with in-house code ^76^. Changes in the positions of amino acids versus retinal were calculated the projection **d**_aa_(*t*)⋅**r** of the center-of-mass displacement (COMD) **d**_aa_(*t*) vector onto the distance vector **r** to retinal at time zero (Fig. 5G). Since the retinal is the source of energy from photon absorption, the motivation of this type of analysis was to compare the changes in an ensemble of rhodopsin trajectories without needing to align the individual structures within an ensemble. Here **r** is the vector between center-of-masses of an amino acid to that of the retinal at time zero; **R**_aa_(0) and **R**_aa_(*t*) denote the center of mass of the amino acid residue at times 0 and *t,* respectively; and **d**_aa_(*t*) = **R**_aa_(*t*) − **R**_aa_(0) is the COMD of the amino acid residue versus retinal. The corresponding *p*-values were calculated for the COMDs of the amino acids versus retinal from both light versus and dark simulations (see above) enabling detection of sub-picosecond motions of residues due to light activation.

### DENSS analysis of double-difference scattering with calculated scattering profiles

To enable fitting of the atomic models from MD to the experimental data, we built a custom version of DENSS-PDB2MRC ^31^ to work explicitly with the experimental time-resolved difference profiles, unlike the conventional tools like CRYSOL ^77^ and DENSS-PDB2MRC which fit standard, absolute scattering profiles, not time-resolved difference scattering profiles. For each of the 120 time points, DENSS-PDB2MRC was used to calculate scattering profiles from the pumped and dark PDBs from a single, randomly chosen molecular dynamics trajectory. The profiles were optimized by using two parameters, namely the bulk solvent density/excluded volume parameter (*c*1) and hydration shell contrast (*c*2) (both in electron density units; e^−^/Å^3^ ) in the *q*-range from 0.06 Å^−1^ to 0.40 Å^−1^ ^31^. For each difference scattering profile, we thus required two independent fitting parameters, (*c*1, *c*2) for the light (pumped) state, and three additional fixed parameters (*c*1 and *c*2 for the dark reference state and a global scale factor). Excellent fits were obtained between the experimental double-difference profiles and the calculated profiles (Fig 5A).

The analysis involved finding the above fitting parameters for both light-activated and dark scattering profiles. We first conducted a dark parameter optimization at two positive time delays (1.0, and 8.0 ps) by minimizing the χ^2^ value between the calculated and experimental rhodopsin minus opsin scattering profiles and simultaneously optimized all six fitting parameters (two for dark, two for each light state), plus one additional global scale factor. An in-house script with iterative optimization using the Nelder-Mead method provided the best dark parameters (i.e. the scale factor and c1/c2 for the dark profiles at these time points) with initial guesses for c1 and c2 used based on electron density of the pure water at room temperature (*c*1 = 0.334 e^−^/Å^3^) and assuming a 10% change in the hydration shell contrast as compared to that of pure water (*c*2 as 0.03 e^−^/Å^3^). Once the dark state parameters were optimized, the two light (pumped) parameters were then optimized for each experimental TR-XSS difference profile independently, while holding the two dark parameters and scale factor fixed for each of the time delays shown in Figure 5B. These parameters are similar to that used in other implicit solvation-based programs used to calculate the scattering profiles such as CRYSOL ^77^. Notably, while any time points in principle can be used to find the dark parameters, we used these time points as they constitute the largest intensities in the double-difference scattering profiles and have relatively higher signal-to-noise ratio as compared to earlier time points.

## Acknowledgments

We dedicate this paper to the memory of Professor John Spence (April 21, 1946–June 28, 2021). Experiments were carried out at the Linac Coherent Light Source (LCLS), a national user facility operated by Stanford University on behalf of the US Department of Energy (DOE), Office of Basic Energy Sciences (OBES). Use of the Linac Coherent Light Source at SLAC National Accelerator Laboratory, is supported by the US Department of Energy, Office of Science, Office of Basic Energy Sciences under Contract No. DE-AC02-76SF00515. Research was carried out at LCLS during beam time LU18, with additional studies during beam times LE71, LM59, LN60, LP99, and LW75. We acknowledge support from US National Institutes of Health awards R01GM133998 (T.D.G.), R01EY012049 (M.F.B.), and R01EY026041 (M.F.B.); the US National Science Foundation awards DBI CAREER 1943448 (R.A.K.), DBI 1565180 (R.A.K. and N.A.Z.), CHE 1904125 (M.F.B.), and MCB 1817862 (M.F.B., T.D.G., P.F., A.G., R.A.K., and N.A.Z.); and the Russian Foundation for Basic Research grant 16-04-00494A (A.V.S.). F.P. acknowledges support by the Knut and Alice Wallenberg foundation (grant no. 2023.0052). We thank the US National Science Foundation for providing funding of this work through the BioXFEL Science and Technology Center grant DBI 1231306 (T.D.G., P.F., R.A.K. and M.F.B.). S.M.D.C.P.. was supported by a Technology Research Initiative Fund (TRIF) predoctoral fellowship from the University of Arizona; L.S.-E. was supported by a Goodman dissertation fellowship; and S.D.E.F. was partially supported by a Goldwater research scholarship. We thank M. T. Marty for the native mass spectrometry studies. The authors are grateful to the University of Arizona Department of Chemistry and Biochemistry Electronics Shop for design and fabrication of experimental apparatus and for support from the Biodesign Center for Applied Structural Discovery at Arizona State University. We thank S. Botha, D. Deponte, M. Holl, C. Kupitz, C. Li, J. Martin-Garcia, and G. Nelson for support with the sample injection and data acquisition. We thank the Center of Integrated Research Computing at the University of Rochester for the provision of computational resources. This research used resources of the Oak Ridge Leadership Computing Facility, which is a DOE Office of Science User Facility supported under Contract DE-AC05-00OR22725. We gratefully acknowledge discussions with H. Chapman, D. Matyushov, R. Neutze, M. Olivucci, T. Sakmar, G. Schertler, I. Schlichting, and S. Smith during the course of this work.

## Funding

National Institutes of Health (GM133998 to T.D.G.; EY012049 and EY026041 to M.F.B.)

National Science Foundation (CAREER DBI-1943448 to R.A.K.; DBI-1565180 to R.A.K. and N.A.Z.; CHE-1904125 to M.F.B.; and MCB-1817862 to M.F.B., T.D.G., P.F., A.G., R.A.K., and N.A.Z.)

Russian Foundation for Basic Research (16-04-00494A to A.V.S.)

US National Science Foundation through the BioXFEL Science and Technology Center (DBI 1231306 to T.D.G., P.F., R.A.K., and M.F.B.)

Technology Research Initiative Fund predoctoral fellowship from the University of Arizona (to S.M.D.C.P)

Goodman dissertation fellowship from the University of Rochester (to L.S.-E.) Goldwater research scholarship from the University of Arizona (to S.D.E.F) Knut and Alice Wallenberg Foundation (2023.0052 to F.P.).

### Author contributions

M.F.B. conceived the experiment and T.D.G., S.M.D.C.P., A.V.S., D.M., P.F., R.A.K. and M.F.B. carried out the experimental design. S.M.D.C.P., A.V.S., X.X., S.D.E.F., N.W., and U.C. developed the rhodopsin protocol, prepared, and characterized the rhodopsin-detergent samples. L.S.-E., A.S., and A.G. carried out and analyzed the molecular dynamics simulations. R.A., K.K., S.L. R.N., S.Z., and R.A.K. developed, tested, and operated the gas dynamic virtual nozzle (GDVN) sample delivery system. S.C., S.B., and R.A.K. set up and aligned the optical parametric amplifier for the pump-probe experiments. M.S.H., M.L., M.D.S., and S.B. set up and aligned the beamline, characterized the beam focus, and calibrated the timing tool. T.D.G., S.M.D.C.P., A.V.S., X. X., S.D.E.F., J.C., R.F., K.K., D.M., F.P., S.C., M.S.H., M.L., M.D.S., S.B., D.M., P.F., R.A.K., and M.F.B. carried out the experiments. T.D.G., C.S.K.M., S.C., S.M., K.K., N.A.Z., F.P., D.M., and R.A.K. analyzed the experimental data. T.D.G., C.S.K.M, A.G., P.F., R.A.K., and M.F.B. wrote and edited the manuscript with discussion, review, and contributions from all authors.

### Data and materials availability

All processed experimental data included in the article and/or SM will be released at the University of Arizona Research Data Repository) upon publication. Raw X-ray data will be deposited in the Coherent X-ray Imaging Data Bank (CXIDB) and released upon publication (https://www.cxidb.org/index.html). Python scripts used in the data analysis and reduction will be made available on GitHub upon publication.The LOOS source code is available for download on GitHub (http://github.com/GrossfieldLab/loos).

